# Engineered Age-Mimetic Breast Cancer Models Reveal Differential Drug Responses in Young and Aged Microenvironments

**DOI:** 10.1101/2024.10.06.616903

**Authors:** Jun Yang, Lauren Hawthorne, Sharon Stack, Brian Blagg, Aktar Ali, Pinar Zorlutuna

## Abstract

Aging is one of the most significant risk factors for breast cancer. With the growing interests in the alterations of the aging breast tissue microenvironment, it has been identified that aging is related to tumorigenesis, invasion, and drug resistance. However, current pre-clinical disease models often neglect the impact of aging and sometimes result in worse clinical outcomes. In this study, we utilized aged animal-generated materials to create and validate a novel age-mimetic breast cancer model that generates an aging microenvironment for cells and alters cells towards a phenotype found in the aged environment. Furthermore, we utilized the age-mimetic models for 3D breast cancer invasion assessment and high-throughput screening of over 700 drugs in the FDA-approved drug library. We identified 36 potential effective drug targets and 34 potential drug targets with different drug responses in different age groups, demonstrating the potential of this age-mimetic breast cancer model for further in-depth breast cancer studies and drug development.

## Introduction

Aging is one of the fundamental factors contributing to various diseases, including breast cancer, which is the most prevalent non-skin malignancy in women. Age-specific cancer incidences rise exponentially until age 55, followed by a slower increasing rate afterwards^1^. During the aging process, cellular and molecular changes occur in cells, leading to a cancerous phenotype. While the significance of age-related cellular alterations in cancer has been widely investigated^2,3^, little has been documented on the effects of the aging microenvironment until recent years. Recent researches have started to highlight the impact of aging on the tumor microenvironment^4^ and how aging affects tumor progression^5,6^. Aging results in alterations of multiple aspects of the microenvironment, including biophysical properties in the ECM, biomolecule secretion, and the associated immune responses. These changes lead to significant differences in ECM characteristics and cell behavior, and further regulate breast cancer therapeutic outcomes and recurrence^7^.

Emerging studies have revealed the crucial contributions of the extracellular matrix (ECM) and its components in breast cancer progression and metastasis^4,6^. In addition to providing structural integrity and sustainability^8^, the ECM influences and regulates various cell behaviors, including survival, proliferation, differentiation^9^, invasion^10^, and immune response^11^ in the breast tissue. These regulations are dictated by the physical characteristics, including fiber structure^12^ and stiffness^13^, and the biochemical compositions, including the structural proteins like collagen^14,15^ and hyaluronic acid^16^, and the soluble factors, like matrix metalloproteinases (MMPs), growth factors, cytokines^17,18,19,20^, and microRNAs (miRNAs)^18,21^. While recent studies have shed light on the impact of age-related changes within breast tissue microenvironment on breast cancer progression and invasiveness^4,6^, much remains unknown about the aging breast tissue microenvironment, hindering the ability to provide guidance for clinical treatments and the development of novel therapeutics.

Although the tumor microenvironment (TME) is extremely complex, it is possible to recapitulate at least the basic components that exert significant influences in tumor progression for in-depth mechanistic studies and drug development. Currently, preclinical studies in cancer drug development have been mostly based on the *in vitro* cytotoxicity and efficacy assays in monolayer cell culture models^22^, which do not recapitulate the three-dimensional (3D) TME, and thus fail to reflect the actual response of tumors in the body to these drugs. Currently, both xenograft and syngeneic *in vivo* models are used in breast cancer drug discovery delivery studies. However, *in* vivo animal models often mismatch the physiological conditions in patients in pathological and aging process. Also, *in vivo* animal studies lack time and cost efficiency and may involve ethical problems when conducted on a larger scale. This discrepancy highly contributes to the inefficient translation of preclinical findings, where 95% of the drugs that are effective in preclinical trials have proven ineffective in clinic^23^, and only 7.5% of drugs tested in Phase 1 trials eventually get approval for clinical use^14^. 3D breast cancer models mimicking the ECM composition, organization, and mechanical properties have been developed over the past years with increasing knowledge of the breast microenvironment^22^. In a study, tumor cells in 3D culture were reported to be less resistant to tirapazamine, a cytotoxic drug in hypoxic conditions, than those in monolayer cell culture^24^. In another study, gene expression profiles of patient samples were analyzed in a 3D in vitro model, and different genes were identified for a better prognosis in ER- and ER+ cancers^25^. The development of 3D *in vitro* models allows drug discovery to better validate the targets *in vitro* and minimize the excessive usage of *in vivo* animal models.

Because of the differences in the aging microenvironment, age also poses a significant impact on drug responses. However, current breast cancer models, including most animal models lack age relevance and fail to predict drug responses in patients from different age groups. Statistically, breast cancer patients over 65 present poorer outcomes with lower survival rates compared to their younger counterparts^26^. Elderly patients are considered underdiagnosed and undertreated. In large randomized clinical trials, elderly patients are usually underrepresented due to accessibility and complication risks. While it is difficult to include more elderly patients in clinical trials, it is difficult to predict clinical outcomes of novel therapies in elderly patients despite the largely affected population. It has been previously identified that drug resistance is relevant to cellular senescence and therapeutics targeting senescence pathways have been under development as adjuvants^27,28^. Recently, with the development of genome profiling, a revelation of drug resistance genes has been an emerging topic. Researchers have started to identify their age and sex-related expression levels and construct computational risk models for predicting patient survival and responses with selected age-related genes^29^. Meanwhile, age-related breast cancer models are urgently needed to assist novel drug screening by identifying drug targets that are more effective in specific age groups, which may require specific attention in further development, trials, and clinical usage.

While the age-related alterations of the TME are not yet fully understood, the fabrication of age-mimetic models utilizing materials with aging characteristics might be favorable for optimizing the fabrication of age-mimetic models. Previously, age-related alterations have been reported in mouse tail tendon collagen^30^, making it a promising candidate for modeling the aging TME. In this study, we investigated the similarity between the age-related properties in mouse breast tissue collagen and tail tendon collagen and assessed the suitability of the tail tendon collagen for generating 3D age-mimetic breast cancer models for breast cancer research and high-throughput drug screening. With isolated collagen with age-resembling characteristics, we constructed a novel age-mimetic breast cancer model with accessible materials that generates an aging microenvironment for cells and alters cells towards a phenotype found in the aged environment. Furthermore, we utilized the age-mimetic models for 3D breast cancer invasion assessment and screened 720 FDA-approved compounds with automated high-throughput drug screening, identifying over 30 drug targets with high efficacy and/or potentially age-altered drug responses in both cell viability and 3D migration. This study substantiated the potential of utilizing animal-generated materials to construct disease models for high-throughput drug screening to include increased patient heterogeneity, facilitating the pre-clinical studies for novel drug development more efficiently.

## Materials and Methods

### Animals

Female mouse tail tendons were used for characterization and collagen isolation in this study. Tissues were harvested from 2–6 months (young) or 20–24 months old (aged) C57BL/6J mice according to the IACUC guidelines (protocol number: 18-05-4687) with the approval of the University of Notre Dame, which has an approved Assurance of Compliance on file with the National Institutes of Health, Office of Laboratory Animal Welfare. Mice were sacrificed in CO_2_ chambers, and tissues were collected and used immediately, or wrapped in aluminum foil, flash frozen in liquid nitrogen, and stored at−80 °C until use.

Mouse 4^th^ mammary gland fat pads were dissected through making one skin incision across the abdomen and a second incision down the midline. The skin was peeled back, and the mammary gland was removed from the skin carefully with small scissors. Mouse tails were collected with small scissors.

### Collagen Isolation

Mouse tail tendon collagen was isolated with a modified established protocol^31^. Isolated tail tendons were stirred in 0.02 N acetic acid in 50 mL tubes on a magnetic stirrer at 4 °C for 72 hours. Solubilized type I collagen was separated through centrifugation at 4,000 × g at 4°C for 20 minutes. Collected supernatant was lyophilized for 48 hours. Lyophilized collagen was weighed and resolubilized in 5 mg/mL in 0.02 N acetic acid on a magnetic stirrer at 4 °C for 72 hours.

### Collagen Characterization

Mouse breast tissues and tail tendons were decellularized with 0.1% SDS for 48 h at 4 °C and delipidized with IPA for 48 h at 4 °C. Morphology of the collagen fibers in aged and young mouse breast tissues and tail tendons were visualized through second harmonic imaging (SHG) with a two-photon microscope (Leica Stellaris 8 Dive). SHG imaging was performed at 25x magnification with optical zoom on the specific sites.

Lectin staining of the collagen fibers were performed on 300 μm cryosectioned native tissues fixed with FITC conjugated triticum vulgare lectin (EY Laboratories) according to manufacturer’s instructions.

### 3D Collagen-Based Breast Tissue Model Preparation

Collagen gels were prepared in 3 mg/mL according to the previously established protocol^31^. Collagen samples were diluted to 3 mg/mL with transglutaminase solution, mixed with 10x PBS, and brought to alkalinity with 0.5 N NaOH on ice. Collagen mixtures were prepared in cylinder-shaped PDMS molds with a 6 mm diameter or in 96-well plates and incubated at 37 °C for 1 h to gel.

### Mechanical Properties

Hydrogel and native breast tissue stiffness was assessed with a Chiaro nanoindenter (Optics 11 Life) with probes with 0.025 N/m stiffness and 10 μm tip radius.

Hydrogel dynamic mechanical properties were evaluated with a TA Instruments HR-2 rheometer fitted with a Peltier stage set to 37 °C. All measurements were performed using a 25 mm parallel plate geometry. Oscillatory strain amplitude sweep measurements were first conducted at a frequency of 20 rad/s. Oscillatory frequency sweep measurements were then conducted at 3% strain after verification that this was in the linear viscoelastic region for the materials.

### SEM

SEM imaging was conducted on the age-mimetic breast tissue models using a gold palladium coating of 9 nm on the samples. Coated samples were imaged at NDIIF with a Magellan 400-field emission scanning electron microscope (FESEM).

### Degradation Study

Collagen gels were prepared and incubated in PBS at 37°C for 7 days to monitor gel degradation. Supernatants and collagen gel samples were collected on day 0, 3 and 7. Collagen concentration released in the supernatants and remained in the collagen gels were measured with a Abcam Total Collagen Assay according to manufacturer’s instructions.

### Cell Culture

Cancerous and normal human mammary epithelial cell lines are used in this study. Green fluorescent protein (GFP)-reporting MDA-MB-231 mammary carcinoma cell line was cultured in cancer cell growth medium (DMEM [high glucose] medium supplemented with 10% FBS [Thermo Fisher Scientific] and 1% penicillin/streptomycin [Corning]). KTB21 human mammary basal epithelial cell line was cultured in epithelial cell growth medium (DMEM [low glucose]: Ham’s F12 [1:3] medium supplemented with 5% FBS [Thermo Fisher Scientific], 0.4 μL/mL hydrocortisone [Sigma], 1% penicillin/streptomycin [Corning], 5 μg/mL insulin [Sigma], 10 ng/mL EGF [Millipore], 6 mg/mL Adenine [Sigma], and 10 mM ROCK inhibitor [Y-27632] [Enzo Life Sciences]). Mouse Mammary Fibroblasts were cultured in a Fibroblast Basal Medium with supplements provided by the manufacturer. Cells were maintained in culture until 90% confluency in a CO_2_ incubator at 37°C and 95% humidity. Cells were passaged using 0.25% trypsin-EDTA, reconstituted in cell growth media, and seeded in culture flasks or plates.

### Cell Proliferation

Cell proliferation was assessed through alamarBlue metabolic assays over the period of 7 days according to manufacturer’s instructions. After culturing for 7 days, epithelial cells were fixed and stained for Ki67. Ki67+ cells and ratios were analyzed with ImageJ.

### Cell Motility

The effect of age-related alterations of collagen on cell motility was assessed through an in-vitro migration assay. Aged and young collagen solutions (3 mg/mL, pH > 7) were prepared and applied to PDMS molds placed in glass bottom culture dishes, and incubated at 37°C for an hour for gelation. The MDA-MB-231 cells and the CellTracker Green CMFDA (Thermo Fisher Scientific)-stained KTB21 cells were washed, treated with trypsin, and reconstituted in culture media at 1.67 million cells/mL. A 30 μL aliquot of the cell suspension was seeded on top of the collagen gels (total of 5,000 cells), and incubated overnight at 37°C for the cells to attach. After cell attachment, fresh media was added, and cells were subjected to time-lapse imaging under an inverted fluorescence microscope for 4 h with 15 min intervals. Each group was performed with two biological replicates. Cells were tracked with Fiji software (NIH).

### Cell Invasion Through Collagen Gels

Effects of the age-related alterations of collagen on breast cancer invasiveness in 3D cell culture were also assessed in vitro with transwell invasion assays. KTB21 cells were labeled with CellTracker Green CMFDA before reseeding for visualization. Transwell inserts were coated with 60 μL of aged or young collagen solution (3 mg/mL, pH > 7) and incubated at 37°C for an hour for gelation. Epithelial cells were seeded onto collagen gels at 30,000 cells per well (n=4). The bottom chambers contained cell medium with an additional 10% FBS as chemoattractant. After incubation for 72 h at 37°C, the transwell inserts were removed and the invaded cells in the bottom chambers were counted using a microscope.

### Age-Mimetic 3D Breast Cancer Model Preparation and Visualization

Tumor spheroids were prepared with Nunclon Sphera 96-well ultra-low attachment plates with GFP-tagged MDA-MB-231 cells in 10,000 cells/well. Cancer cells were seeded and cultured for 48 h to form spheroids. Formed spheroids were encapsulated in 40 μL of previously described 3 mg/mL isolated aged or young mouse tail tendon collagen solutions and incubated at 37 °C for 1 h to gel.

3D age-mimetic breast cancer models are cultured in DMEM complete medium as described for 2 days and fixed with 4% PFA. Fixed models were stained for type I collagen (Abcam ab260043) and DAPI and visualized microscopically.

### Automated 3D Breast Cancer Model Monitoring

Automated monitoring of the 3D models was performed with a BioTek BioSpa 8 Automated Incubator coupled with a BioTek Cytation 5 Cell Imaging Multimode Reader. Age-mimetic 3D breast cancer models were imaged on day 0,1,2,3,5,7 with automated spheroid size measurements to monitor spheroid growth or shrinkage. Start and endpoint images were also further analyzed with ImageJ to quantify 3D cell invasion.

### High-Throughput Drug Screening

High-throughput drug screening was performed on plate 1-9 from an FDA-approved drug library (Selleck L1300) with the 3D age-mimetic breast cancer models and a Biomek i7 Automated Workstation. Models were cultured overnight after spheroid encapsulation and drug addition (10 μM) was performed on day 1 with an endpoint CellTiter-Glo 3D Cell Viability Assay on day 5. Automated imaging was also performed on day 1 immediately after drug addition and day 5 before endpoint analysis.

### Selected Drug Efficacy

Selected drug targets were prepared for dose-dependent assessment with a serial dilution from 2.5 nM to 150 μM. IC50 assessment was performed with the 3D age-mimetic breast cancer models and a Biomek i7 Automated Workstation in a PrimeSurface 384U 3D cell culture plate. Models were cultured overnight after spheroid encapsulation and drug addition was performed on day 1 with an endpoint CellTiter-Glo 3D Cell Viability Assay on day 5. IC50 Analysis was performed with Graphpad Prism 10.

### Statistical Analysis

Data were analyzed for statistical significance with Prism 10 (Graphpad). Two-tailed unpaired student’s *t*-test was applied to compare the difference between two groups and one-way ANOVA followed by Tukey’s HSD correction was performed to compare the differences between multiple groups. Outliers were identified using the ROUT method with *Q* = 1% and eliminated. Data are presented as the mean ± standard deviation (SD).

## Result and Discussion

### Aged Tail Tendon Collagen Resembles Collagen Fibers in Aged Breast dECM

SHG imaging of the decellularized mammary gland tissue slices demonstrates the difference between the morphology of the aged and young breast collagen fibers. As discussed in a previous study^6^, while young collagen fibers were more straight and aligned, the aged collagen fibers tend to be curvier. The similar differences were observed in the tail tendons. Although the structure of the tail tendon restricts the collagen fibers to better align along the bones, resulting in the almost perfectly straight and aligned morphology in the young tails, fibers in the aged mouse tail tendon still presented significant curvature. Our previous studies have demonstrated that collagen scaffolds with curvier fibers promoted mammary epithelial cell invasiveness in the context of obesity^32^. The similar age-related differences in the morphology of the collagen fibers indicated the possibility of utilizing the aged mouse tail tendon collagen to generate breast tissue mimicking 3D bioscaffolds for breast cancer research.

Other than the morphology, researchers^30,33^ have previously reported the age-related alterations in glycation levels in the tail tendon collagen and how glycation alters breast cancer progression and invasion^34^. Previous studies have reported that advanced glycation endproducts would promote breast cancer cell proliferation, migration, and invasion, resulting in a higher likelihood of breast cancer occurrence and metastasis in diabetic patients^33^. Lectin staining further discovered that SNA-I, WGA, and PHA-E are highly expressed on the aged collagen fibers in both types of tissues with low expression in the young groups from both types of tissues. These further confirmed that the aged tail tendon collagen resembles both morphological and biochemical age-related alterations in aged mammary gland tissues, making them a good source of material for generating 3D breast cancer models mimicking the aging breast microenvironment.

### 3D Collagen Based Age-Mimetic Breast Tissue Models Demonstrated Similar Mechanical Properties

Circular dichroism showed that soluble collagen isolated from aged and young mouse tail tendons exhibit analogous secondary structures and almost identical melting points. Onset of the loss of structural integrity occurred in the aged collagen at a slightly lower temperature, indicating the incrementally lower thermal stability of the aged samples.

SEM imaging of the hydrogel generated from the isolated collagens demonstrated that although the aged collagen fibers exhibited significantly higher thickness, the pore sizes of the two groups of hydrogels are remarkably similar. The similarity in the structure of the 3D hydrogel systems would present similar physical barriers and drug diffusion properties to cancerous cells in further breast cancer research.

As for the mechanical properties, nanoindentation of both the breast tissues and collagen gels with Piuma nanoindenter measured Young’s modulus lower than 80 Pa, indicating that both the breast tissues and hydrogels were very soft and the differences in stiffness could be negligible. Rheology assessments of the hydrogel systems showed that both groups of collagen gels were very weak hydrogels, with a storage modulus of 19.00 ± 1.65 Pa for the aged groups and 20.21 ± 1.62 Pa for the young groups, and a loss modulus of around 2.6 Pa for both groups. The proximity of the dynamic strength of the two groups of hydrogels further confirmed that cells are unlikely to undergo different stress due to substrate stiffness in the 3D age-mimetic breast tissue models.

To validate the stability of the hydrogel system for time-dependent drug response studies, degradation of the collagen-based hydrogels was assessed with total collagen assays. Both groups of collagen hydrogels maintained over 90% of the collagen contents within the 7-day incubation period, with around 5 μg of collagen released into the buffer system in 3 days and less than 10 μg of collagen released total in 7 days.

All hydrogels were able to maintain integrity after 7-day incubation, demonstrating the model’s suitability for long-term drug response monitor.

### Cells Acquired Age-Related Properties from Age-Mimetic Breast Tissue Models

To assess the cell viability when cultured in the age-mimetic breast tissue models, alamarBlue assay showed that in both aged and young models, both cancerous and normal epithelial cells maintained similar metabolic rates throughout the culture period of 7 days. Ki67 staining further demonstrated that both cancerous and normal epithelial cells had maintained similar proliferative ability after being cultured in the aged and young model hydrogel systems.

PCR analysis of both epithelial cells (Figure 3C) cultured in the age-mimetic breast tissue models demonstrated that cells cultured in the aged models expressed significantly higher levels of SERPINE1, an acknowledged aging marker, similar to cells cultured on the aged dECM. Upon further investigation, the same molecular change was also observed in mouse mammary fibroblasts (Supplementary Figure S1), the main regulators of mammary gland microenvironment. The upregulation of aging markers in mammary stromal cells, including epithelial cells and fibroblasts in the aged breast tissue models further verified that cells were able to assimilate age-related alterations on the molecular level.

**Figure 1.**
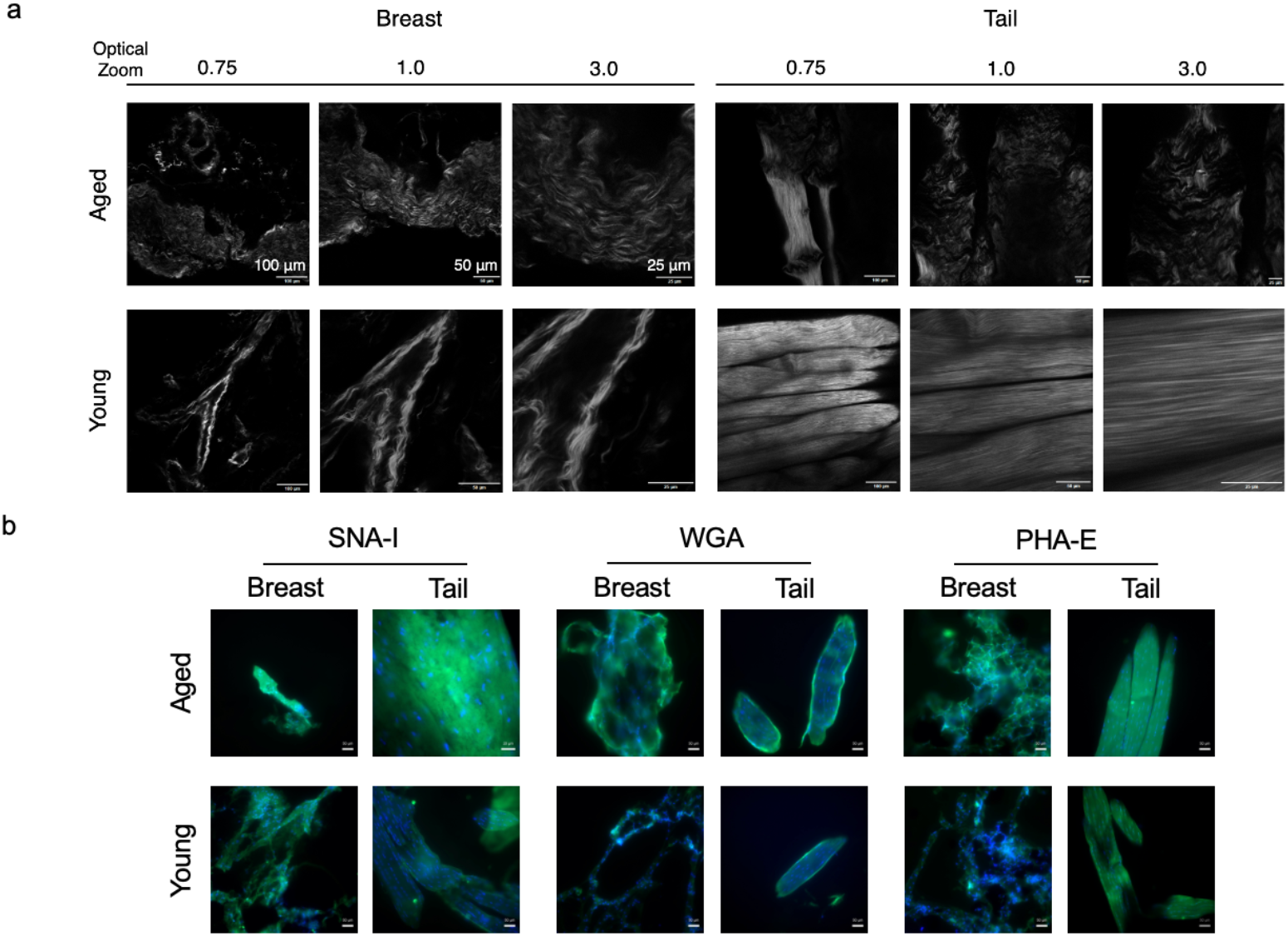
Characterization and comparison of collagen fibers in native aged and young mouse breast tissue sections and tail tendons. A) Second harmonic images of decellularized aged and young mouse 4^th^ mammary gland tissue and tail tendon. B) Lectin staining of decellularized aged and young mouse 4^th^mammary gland tissue and tail tendon sections.

**Figure 2.**
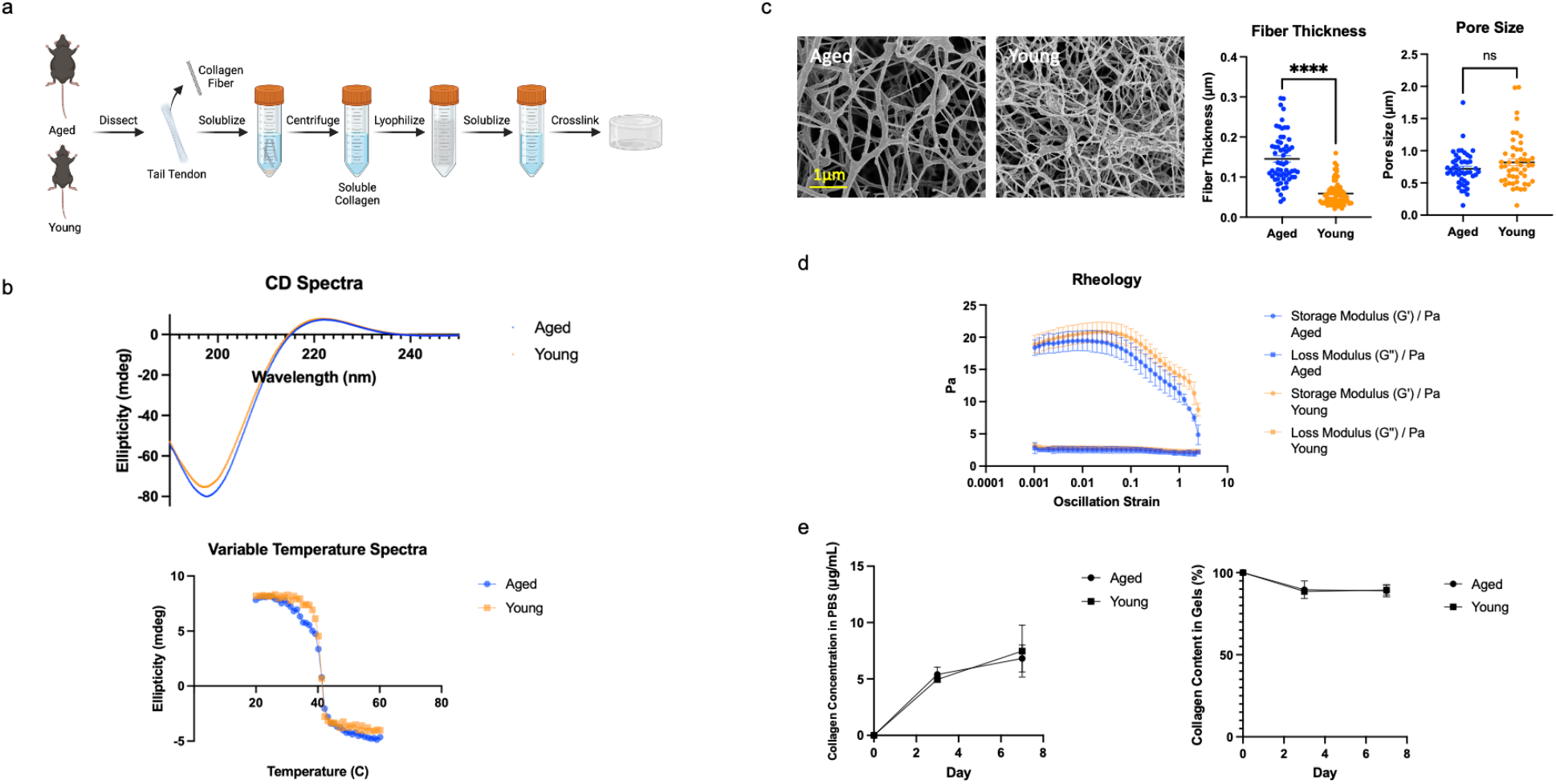
Characterization of isolated soluble collagen and constructed collagen-based hydrogels from aged and young mouse tail tendon. A) Scheme of soluble mouse tail tendon collagen isolation. B) Circular dichroism of isolated aged and young soluble tail tendon collagen (n=2). C) SEM images of 3D age-mimetic breast tissue models and quantification of collagen fiber thickness and hydrogel pore sizes (n=3). D) Dynamic rheology of 3D age-mimetic breast tissue models (n=3). E) Degradation of collagen-based 3D age-mimetic breast tissue models over 7 days incubation (n=4). Data presented as the mean ± SD. ANOVA followed by Tukey’s post hoc was applied for statistical significance. *p<0.05, **p<0.01, ***p<0.001, and ****p<0.0001.

**Figure 3.**
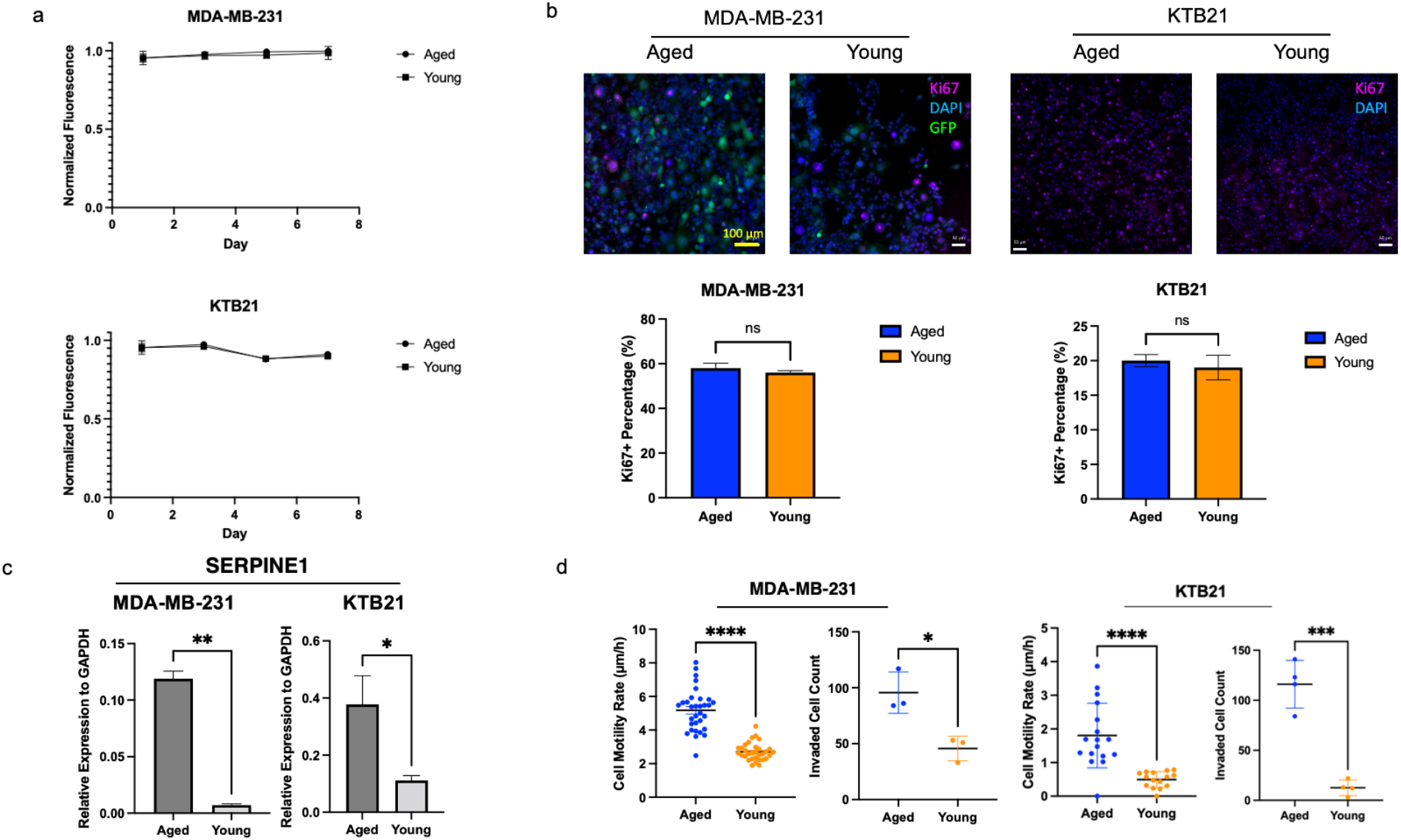
Assessments of cellular behaviors in the 3D age-mimetic models. A) Cell metabolism epithelial cells cultured in 3D age-mimetic breast tissue models monitored with alamarBlue assays (n=3). B) Ki67 proliferation marker staining and quantification (n=3). C) Relative expression of aging marker SERPINE1 in mammary epithelial cells and fibroblasts cultured in 3D age-mimetic breast tissue models for 7 days (n=4). D) Epithelial cell motility (n=3) and transwell invasion (n=4) of epithelial cells cultured in 3D age-mimetic breast tissue models. Data presented as the mean ± SD. ANOVA followed by Tukey’s post hoc was applied for statistical significance. *p<0.05, **p<0.01, ***p<0.001, and ****p<0.0001.

Cell motility tests with both cancerous and normal epithelial cells seeded on aged and young breast tissue models demonstrated that cells presented a significantly higher motility rate on the aged models compared to the young models (p<0.0001 for both cell types). Similarly, epithelial cells also showed significant higher invasiveness through the aged models (p= 0.016 for MDA-MB-231 cells and p=0.0002 for KTB21 cells). Both cell types presented similar age-related behavioral differences as cells cultured on aged and young dECM in our previous study^6^. As epithelial cells presented similar viability and proliferation in the aged and young models, their cellular motility and invasion differences were not likely to be a result of different cell viability. Rather, epithelial cells in the aged breast tissue models undergo cellular alterations due to the differences in the collagen properties and transform into a more invasive phenotype. Cellular behaviors indicated that epithelial cells were able to acclimate to the aged or young microenvironment generated in the 3D age-mimetic models and translated the environmental differences into different cellular responses, recapitulating the cell behaviors in the aging microenvironment.

Given that the mechanical properties of the two aged and young collagen-based hydrogels have been proven to be similar in the two groups of age-mimetic models, these alterations in cellular activities are likely to be induced by the differences of the collagen fibers in animals from different age groups. The 3D age-mimetic breast tissue models generated bioscaffolds mimicking the aging microenvironment and cultivated the cells to present more aging-associated characteristics.

### 3D Tumor Spheroids Demonstrated More Invasive Behavior in Age-Mimetic Breast Cancer Models

As shown in Figure 4a, we successfully generated tumor spheroids and embedded them in age-mimetic breast tissue models with isolated tail tendon collagen to construct the 3D age-mimetic breast cancer models. Tumor spheroids were able to maintain structural integrity after collagen gel encapsulation with the collagen fibers covering the spheroids.

**Figure 4.**
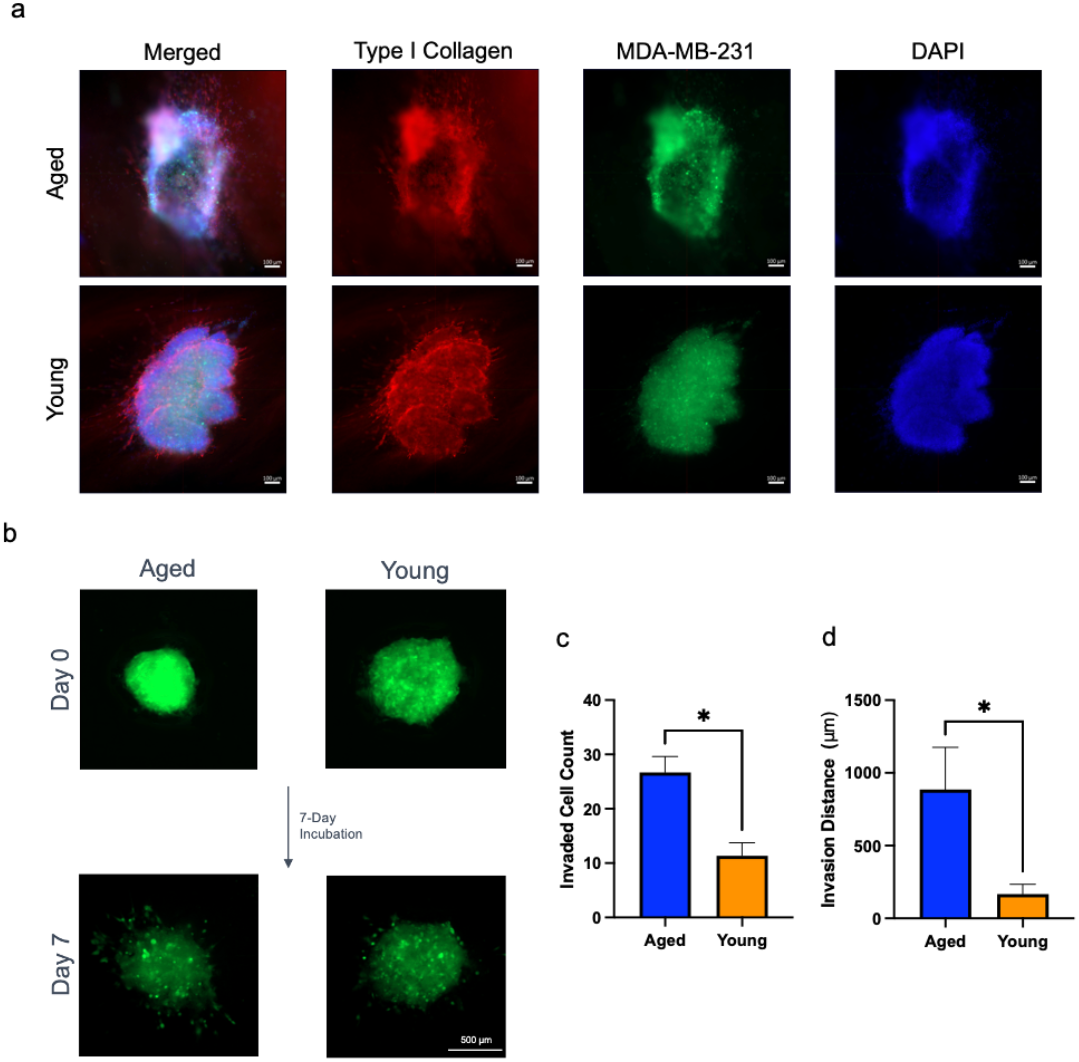
Automated monitoring of 3D cell invasion in age-mimetic breast cancer models. A) Immunofluorescence imaging of 3D age-mimetic breast cancer models with GFP-tagged MDA-MB-231 cells stained with DAPI and bioscaffolds stained for type I collagen B) Automated imaging monitored MDA-MB-231 cell 3D invasion over a period of 7 days. C) Quantification of invaded cell counts from 3D age-mimetic breast cancer models (n=6). D) Invasion distance of 3D invasion within the 3D age-mimetic breast cancer models, analyzed by Image J (n=6). Data presented as the mean ± SD. ANOVA followed by Tukey’s post hoc was applied for statistical significance. *p<0.05, **p<0.01, ***p<0.001, and ****p<0.0001.

With the automated imaging systems, we were able to visualize 3D cell migration from the spheroids over the period of 7 days incubation. Quantification of the 3D cell migration showed significantly higher invaded cell count (p=0.03) as well as invasion distances (p=0.016) in the aged breast cancer model group, in correspondence with the cell behaviors in 2D experiments, where tumor spheroids spread further and faster on the aged dECM. The 3D migration experiments further validated that the 3D age-mimetic breast cancer models were able to recapitulate the invasion behaviors in the aged and young breast tumor microenvironment.

### High-Throughput Screening Identified Drugs with Different Responses from Age-Mimetic Breast Cancer Models

High-throughput screening of 720 compounds from the FDA-approved drug library (Figure 5a) identified 36 compounds with over 50% efficacy in reducing cancer cell survival in both aged and young breast cancer models over the period of 5 days, with 27 compounds only efficient in young models and 7 compounds with improved performance on the aged models. We further analyzed the 3D migration of cancer cells in the age-mimetic breast cancer models to identify the compounds significantly limiting the migration of cancer cells within the models (Figure 5b). Among the 36 compounds with over 50% efficacy in both groups, 15 compounds were limiting the cancer cell migration in both groups while 7 compounds were only effective in limiting 3D migration in the young groups.

**Figure 5.**
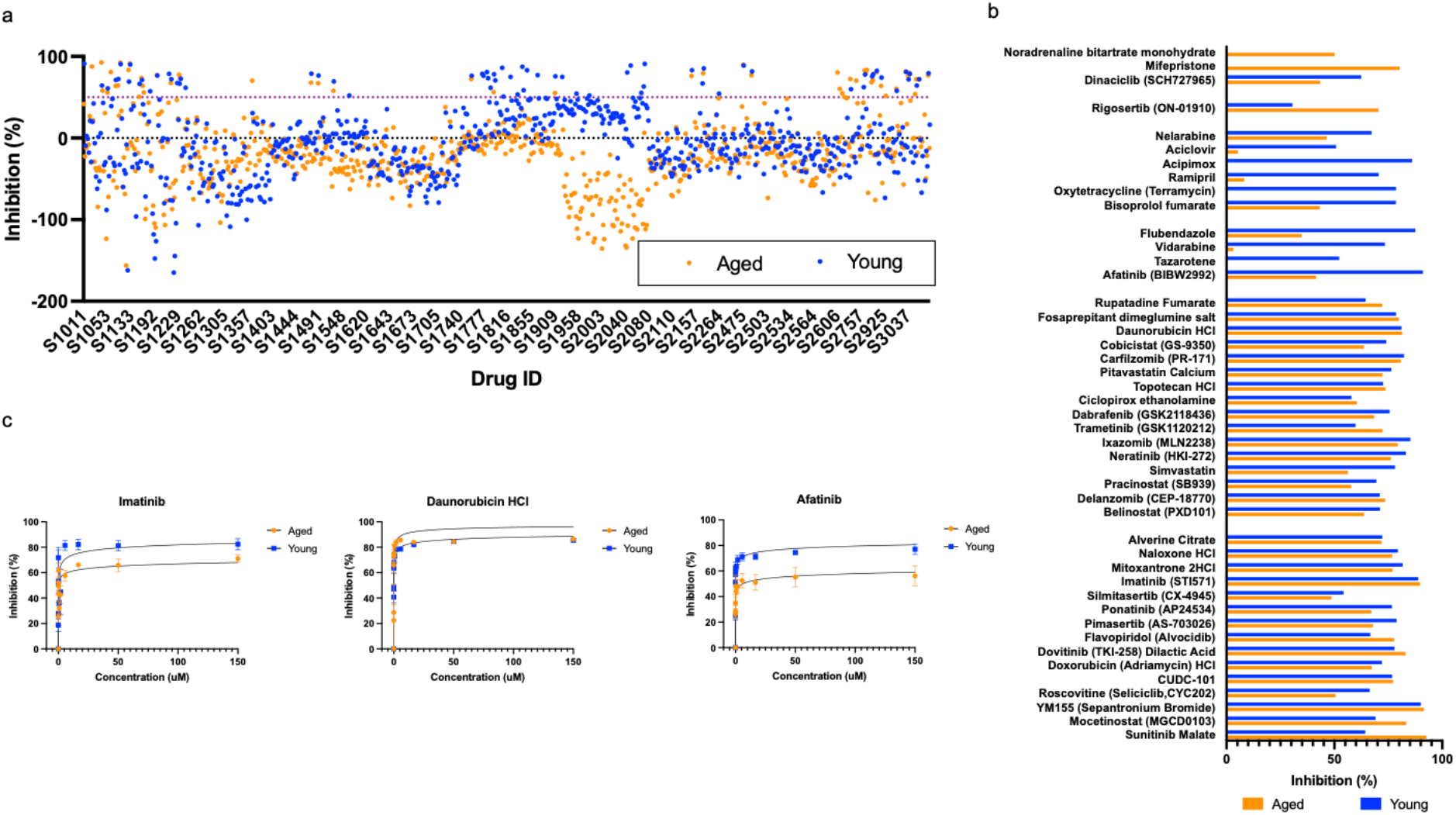
High throughput drug screening with 3D age-mimetic breast cancer models. a) High throughput screening of 720 compounds from the FDA-approved drug library utilizing the 3D age-mimetic breast cancer models. b) Hit selected with > 50% efficacy from the HTS . c) Dose-response effect of the three major lead drug on the 3D age-mimetic breast cancer models (n=4).

To evaluate the dose-dependent responses from the age-mimetic breast cancer models, we selected 3 drugs with over 80% efficacy in the drug library screening experiments, with 2 compounds effective in both groups and 1 compound with significantly higher efficacy in the young group (Figure 5c). Imatinib is an FDA-approved treatment for chronic myeloid leukemia and gastrointestinal stromal tumors through targeting tyrosine kinases^35^, which has been used for metastatic breast cancer treatments^36–39^. Imatinib is a multi-target inhibitor of v-Abl, c-Kit and PDGFR with IC50 of 0.6 μM, 0.1 μM and 0.1 μM, respectively. The dose-dependent assessments demonstrated IC50 of 0.22 μM and 0.10 μM in the 3D aged and young mimicking models, respectively. Daunorubicin HCl inhibits both DNA and RNA synthesis and inhibits DNA synthesis with Ki of 0.02 μM. In current breast cancer studies, this compound is more commonly used combined with other treatments including tamoxifen and amlodipine^40^. It has also shown the potential of reversing drug resistance to TNF-related apoptosis-inducing ligands through downregulation of Mcl-1^41^. In this study, Daunorubicin HCl showed antiproliferative effects in both groups of age-mimetic models with IC50 of 0.01 μM in both groups, showing its high potency in antiproliferation in different age groups. Afatinib irreversibly inhibits multiple checkpoints in the EGFR/HER2 signaling pathway including EGFR(wt), EGFR(L858R), EGFR(L858R/T790M) and HER2 with IC50 of 0.5 nM, 0.4 nM, 10 nM and 14 nM, respectively; It has been shown to be 100-fold more active against Gefitinib-resistant L858R-T790M EGFR mutant. Multiple treatments involving this compound in clinical trials have completed phase II studies^42–44^. However, in the 3D age-mimetic models, Afatinib showed IC50 of 5.6 μM in the aged group and 0.02 μM in the young group, indicating this compound might not be as effective for patients of relatively elder age. Results from this high-throughput screening of the FDA-approved library also identified multiple compounds that have not been studied for breast cancer treatment. These results provide valuable insights for further therapeutic research, paving the way to identifying promising compounds in breast cancer treatment and improving therapeutic strategies for patients in different age groups. The dose-dependent experiments further demonstrated the reproducibility of studies utilizing the age-mimetic breast cancer models, indicating its potential in novel drug screening in future drug development studies. Furthermore, this age-mimetic model could incorporate co-culture of other cell types from mammary stromal cells to cardiac cells to evaluate the cytotoxicity of the drugs. Clinically, potential cardiotoxicity and other side effects of chemotherapy treatments have been a significant factor of the undertreatment of elderly patients. Utilizing this age-mimetic model in drug development could allow researchers to identify more safe and effective therapeutics for the elderly patients, creating a more accessible and promising plan of chemotherapy.

## Conclusion

In this study, we fabricated a 3D age-mimetic breast cancer model with isolated aged and young type I collagen for high throughput drug screening. This study identified the similarity in age-related alterations in collagen fibers from mouse breast tissues and tail tendons demonstrating the possibility of generating 3D age-mimetic breast cancer models with isolated mouse tail tendon collagen. Collagen fibers in the two tissues share similar age-related alterations in curvature and glycosylation. Mammary stromal cells adopt age-related characteristics in the generated aged scaffolds and demonstrated similar cellular behavioral changes as cells seeded on aged and young dECM, with significant increases in motility and invasiveness in cells cultured in the aged models. The 3D age-mimetic breast cancer models were utilized to demonstrate the increased 3D cell invasion inside the aged ECM model. Moreover, the age-mimetic models were utilized to screen for potential drug targets in an FDA-approved drug library with over 700 compounds and identified potential drug targets with high efficacy or different drug responses in different age groups. This model could be a standard method for efficiently screen novel drug compounds for breast cancer treatment and serve as a useful tool for clinical drug selection for patients in different age groups.

## Acknowledgements

This study is funded by NIH award number 1R01CA275423-01A1.

This study is supported by the Interdisciplinary Interface Training Program (IITP) Grant, made possible by the Walther Cancer Foundation.

We would like to acknowledge Professor Siyuan Zhang for the gift of the MDA-MB-231 cells used in this study and Dr. Harikrishna Nakshatri of Indiana University for the gift of the KTB21 cells used in this study. Mouse tissues were kindly provided by Professor Sharon Stack and Professor Siyuan Zhang.

The authors acknowledge the use of the Electron Microscopy Core of the Notre Dame Integrated Imaging Facility, a designated core of the NIH-funded Indiana Clinical and Translational Sciences Institute. We thank the Materials Characterization Facility (MCF) for the use of the HR-2 Discovery Hybrid Rheometer, TA Instruments. The MCF is supported by Notre Dame Research.

The schematics in some figures were created using BioRender.com.

## Supplementary Figures

**Figure S1.**
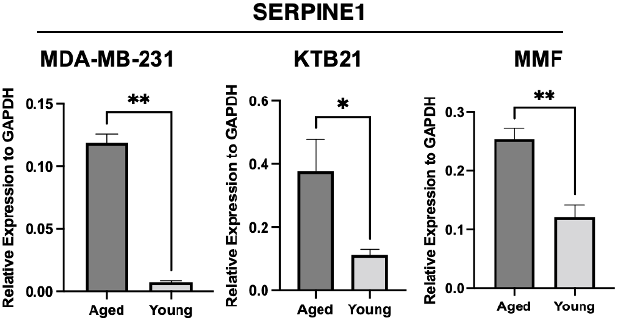
Relative expression of aging marker SERPINE1 in mammary epithelial cells and mammary fibroblasts cultured in 3D age-mimetic breast tissue models for 7 days (n=4). Data presented as the mean ± SD. ANOVA followed by Tukey’s post hoc was applied for statistical significance. *p<0.05, **p<0.01, ***p<0.001, and ****p<0.0001.

